# Distributed Biotin-Streptavidin Transcription Roadblocks for Mapping Cotranscriptional RNA Folding

**DOI:** 10.1101/100073

**Authors:** Eric J. Strobel, Kyle E. Watters, Julius B. Lucks

## Abstract

RNA molecules fold cotranscriptionally as they emerge from RNA polymerase. Cotranscriptional folding is an important process for proper RNA structure formation as the order of folding can determine an RNA molecule’s structure, and thus its functional properties. Despite its fundamental importance, the experimental study of RNA cotranscriptional folding has been limited by the lack of easily approachable methods that can interrogate nascent RNA structures at nucleotide resolution during transcription. We previously developed cotranscriptional selective 2’-hydroxyl acylation analyzed by primer extension sequencing (SHAPE-seq) to simultaneously probe all of the intermediate structures an RNA molecule transitions through during transcription elongation. Here, we improve the broad applicability of cotranscriptional SHAPE-Seq by developing a sequence-independent streptavidin roadblocking strategy to simplify the preparation of roadblocking transcription templates. We determine the fundamental properties of streptavidin roadblocks and show that randomly distributed streptavidin roadblocks can be used in cotranscriptional SHAPE-Seq experiments to measure the *Bacillus cereus crcB* fluoride riboswitch folding pathway. Comparison of EcoRI_E111Q_ and streptavidin roadblocks in cotranscriptional SHAPE-Seq data shows that both strategies identify the same RNA structural transitions related to the riboswitch decision-making process. Finally, we propose guidelines to leverage the complementary strengths of each transcription roadblock for use in studying cotranscriptional folding.

## Introduction

The capacity for RNA to fold into sophisticated structures is integral to its roles in diverse cellular processes including gene expression, macromolecular assembly, and RNA splicing (1,2). Because RNA folding can occur on a shorter timescale than nucleotide addition by RNA polymerase (RNAP) (3-5), a nascent RNA can transition through multiple intermediate structural states as it is synthesized (6). Pioneering experimental studies of RNA cotranscriptional folding used biochemical methods to characterize these RNA structural intermediates (6-8) and more recently, single-molecule force spectroscopy has been used to directly observe RNA folding during transcription by measuring changes in RNA extension in real time (9). However, the lack of a robust method to directly interrogate RNA structure at nucleotide resolution during transcription has so far limited our ability to fully investigate the fundamental principles of RNA cotranscriptional folding and its impact on generating functional RNA structural states that govern fundamental biological processes.

We recently addressed this technological gap by developing cotranscriptional SHAPE-Seq to measure nascent RNA structures at nucleotide resolution (10). SHAPE-Seq combines chemical RNA structure probing with high-throughput sequencing to simultaneously characterize the structure of RNAs in a mixture (11-13). Chemical modification of a target RNA is accomplished using any of the suite of SHAPE probes available that react with the RNA 2’-hydroxyl at ‘flexible’ regions of the molecule, such as unpaired nucleotides in single stranded regions and loops (14). After reverse transcription (RT), modified nucleotides can be detected as truncated RT products using high-throughput sequencing. The resulting sequencing reads are then used to generate a ‘reactivity’ value for each nucleotide in each RNA (15). SHAPE-Seq reactivities represent the relative flexibility of each nucleotide of an RNA: highly reactive nucleotides tend to be single-stranded, whereas nucleotides with low reactivities tend to be constrained by base-pairing or other intra- or intermolecular interactions (11,16).

Cotranscriptional SHAPE-Seq combines the ability of SHAPE-Seq to characterize complex mixtures of RNAs with *in vitro* transcription in order to simultaneously interrogate the structure of all intermediate lengths of a nascent target RNA. Each intermediate length is probed in the context of stalled transcription elongation complexes (TECs) (10) which are generated by constructing a DNA template library containing a promoter, a variable length of the target RNA template, and an EcoRI site. Within 30s of the start of *in vitro* transcription, TECs are blocked by a catalytically dead EcoRI_E111Q_ mutant (Gln111) (17,18) bound to the EcoRI sites (10) and treated with either the fast acting SHAPE reagent benzoyl cyanide (BzCN, t_1/2_ of 250 ms) (19) or dimethyl sulfoxide (DMSO) as a control. RNAs are then quickly extracted and processed for paired-end sequencing to identify transcript length and SHAPE-modification position as described previously (11). RNA structural states that persist on the order of seconds are interrogated to provide “snapshots” of kinetically trapped intermediates that reveal key transitions within RNA folding pathways (10)

The cotranscriptional SHAPE-Seq experiment requires that stalled TECs can be generated for all intermediate lengths within a target RNA sequence at once. Gln111 was initially selected as a roadblock because its ability to halt *Escherichia coli* RNAP is both robust and well characterized (18). However, the use of Gln111 comes with a number of drawbacks. Constructing the Gln111 DNA template library requires a unique primer set that encodes every stop for each RNA target and therefore contributes substantially to experimental costs, which is exacerbated during mutational analysis as many additional primers are required in order to preserve the mutation in the DNA template library. Further, intermediate lengths that are poorly amplified create the potential for gaps in the experimental data, which is particularly problematic for highly repetitive sequences. Last, RNA sequences that contain an internal ‘GAATTC’ must be mutated to access structural information downstream or an alternative transcription roadblock (20,21) must be used. Thus, the development of a sequence-independent roadblocking strategy is highly desirable in order to reduce both experimental costs and time, thereby facilitating a broad application of cotranscriptional SHAPE-Seq to the study of how RNA folding directs RNA function.

Here we develop a sequence-independent method for halting TECs at all positions across a DNA template using streptavidin (SAv) as a transcription roadblock and combine this method with SHAPE-Seq to characterize cotranscriptional RNA folding pathways at nucleotide resolution (Fig. 1). We start by characterizing the robustness of SAv transcription roadblocks in the context of *in vitro* transcription. We then implement SAv roadblocking in the cotranscriptional SHAPE-Seq framework using randomly biotinylated DNA templates to capture TECs across all transcript lengths in a general workflow that can be applied to any RNA sequence. A comparison of the SAv and Gln111 roadblocking strategies using the *Bacillus cereus crcB* fluoride riboswitch (22) as a model system allowed us to identify technical distinctions between SAv and Gln111 roadblocking and we propose experimental strategies that leverage the complementary strengths of each approach. The robust and sequence-independent nature of SAv roadblocking is a powerful addition to the cotranscriptional SHAPE-Seq method that uses reagents that are all commercially available, reduces experimental costs, and simplifies materials preparation. Together, these improvements increase the accessibility of cotranscriptional SHAPE-Seq to a broader user base to study cotranscriptional RNA folding.

**Figure 1.**
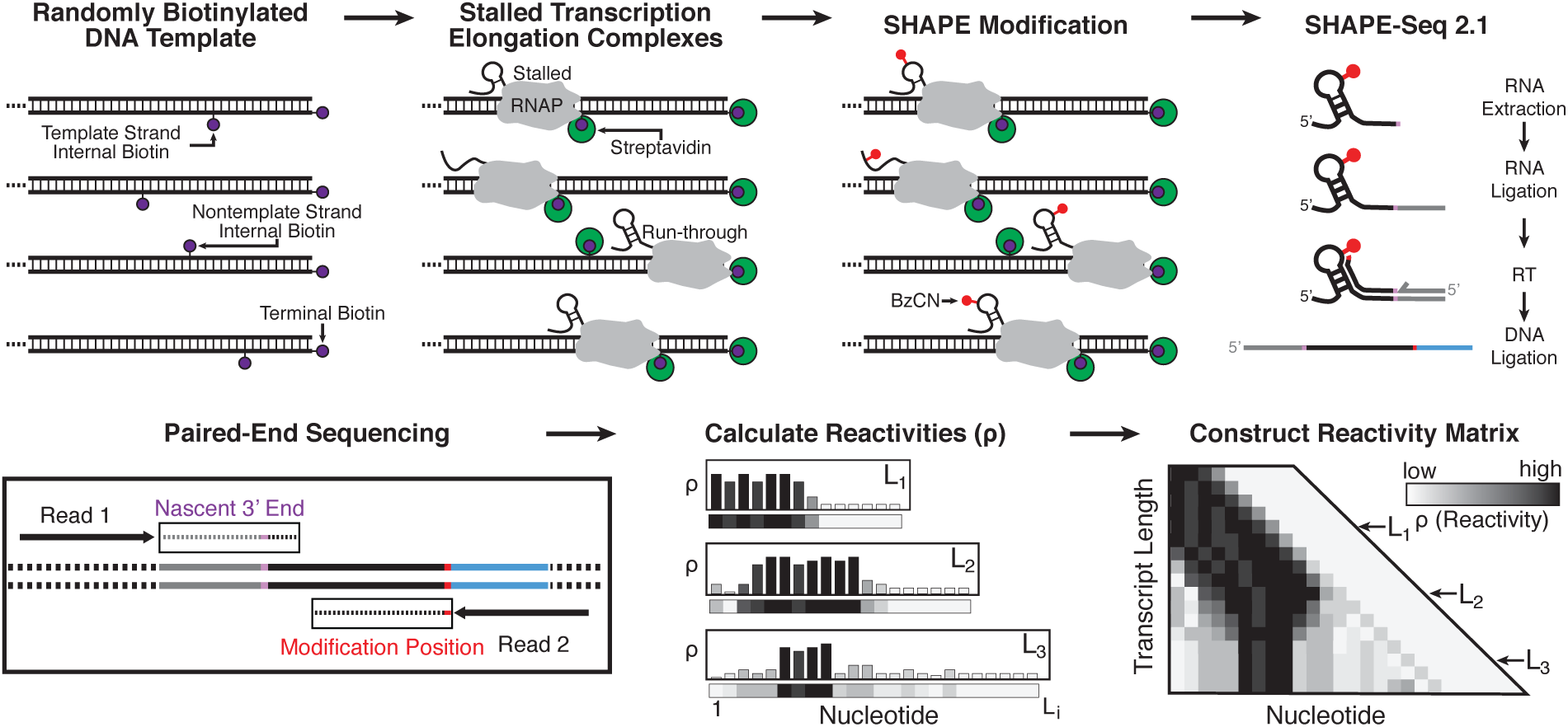
Overview of streptavidin (SAv) roadblocking in cotranscriptional SHAPE-Seq. DNA templates containing random internal biotin modifications and a 5’ terminal biotin modification on the template strand are prepared by PCR amplification. Following open complex formation and incubation with SAv, single-round transcription is initiated to generate stalled transcription elongation complexes. After 30s, nascent RNAs are treated with either BzCN (+) or DMSO (−), extracted, and processed for paired end sequencing using the SHAPE-Seq v2.1 protocol (11). Transcript length and SHAPE modification position are identified by paired-end sequencing and used to calculate a SHAPE reactivity profile for each transcript. Individual reactivity profiles are stacked to generate a reactivity matrix that contains nucleotide resolution structural data about the cotranscriptional folding pathway. The ‘SHAPE-Seq 2.1’, ‘Paired-End Sequencing’, ‘Calculate Reactivities’, and ‘Construct Reactivity Matrix’ panels are adapted from Watters et al., 2016 (10).

## Results

### The efficiency of streptavidin transcription roadblocking is DNA strand dependent

SAv roadblocking (23,24) has previously been shown to prevent E. *coli* RNAP from “running off’ the DNA template during *in vitro* transcription (9). Typically, SAv roadblocks are introduced into a DNA template to prevent run-off transcription by including a 5’-biotinylated reverse primer during template amplification (25). Recently, a terminal biotin-SAv roadblock was used to halt a TEC to facilitate SHAPE probing of a single RNA transcript in complex with RNAP (26). Cotranscriptional SHAPE-Seq, however, requires the distribution of stalled TECs across all DNA template positions. Thus, the use of SAv roadblocking in cotranscriptional SHAPE-Seq requires random biotinylation of the DNA template during PCR, which inherently biotinylates both the template and nontemplate strands of the DNA duplex. Because each DNA strand makes distinct interactions with RNAP, a SAv roadblock in the template strand may not have an equivalent effect on RNAP compared to the nontemplate strand.

To measure the efficiency of template vs. nontemplate strand SAv roadblocks, we performed *in vitro* transcription using DNA templates that were biotinylated at a single internal position downstream of the promoter in the template strand (positions +33 or +42) or in the nontemplate strand (position +33) (Fig. 2A). Template strand SAv roadblocks stall TECs with 80-87% efficiency as a cluster of stops 7-13 nucleotides (nts) upstream of the biotinylation site. In contrast, nontemplate strand SAv roadblocks stall TECs as a more defined stop, but with only 30% efficiency (Fig. 2A). The superior roadblocking efficiency of the template strand SAv roadblocks is consistent with the trajectory of each DNA strand through RNAP: the template strand must traverse the RNAP active site deep within the polymerase internal channel, while the nontemplate strand is less sterically constrained (27). The clustered distribution of TECs stalled by a SAv roadblock is distinct from the defined stop 14 nt upstream of the EcoRI for TECs stalled by Gln111 (18), and is likely due to the flexible linker between biotin and the DNA template.

**Figure 2.**
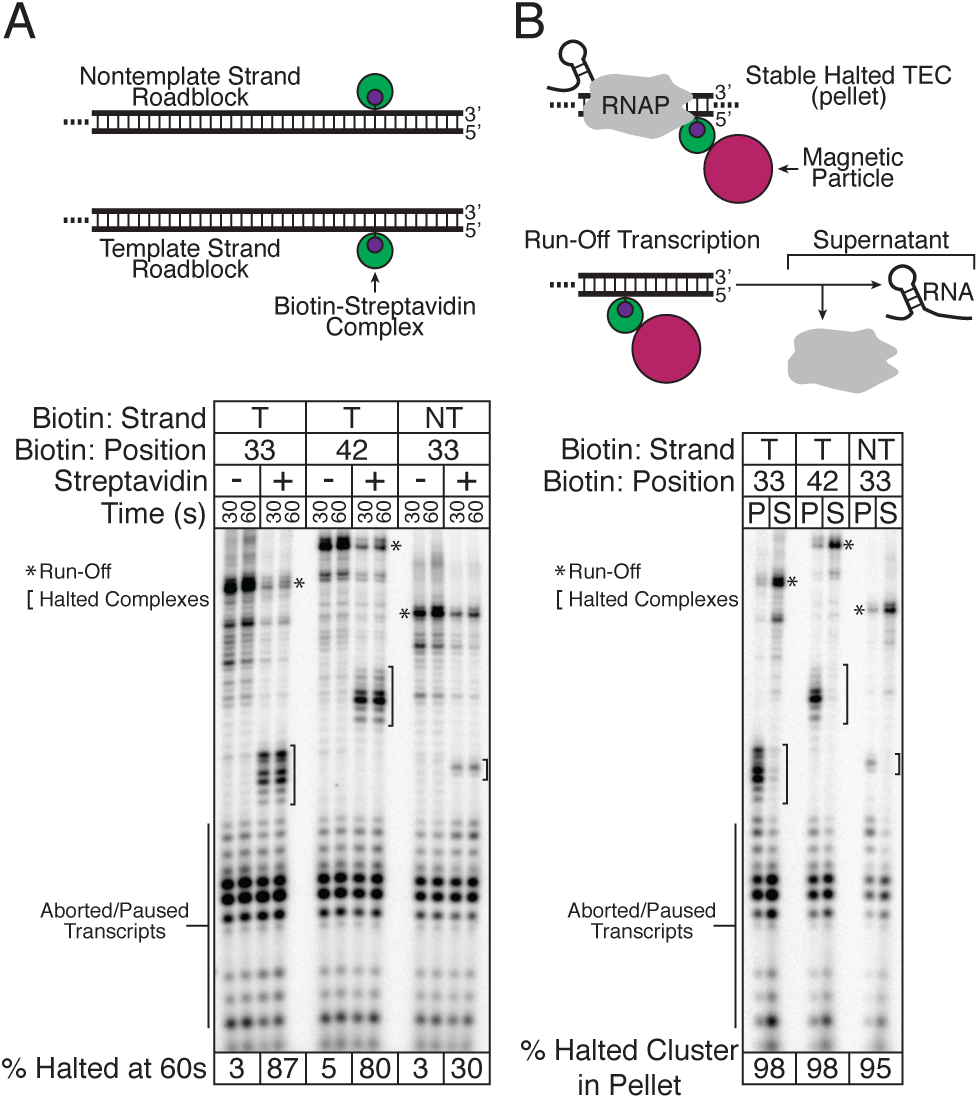
Characterization of SAv transcription roadblocks. **(A)** Template strand (T) SAv roadblocks halt TECs with greater efficiency than nontemplate strand (NT) SAv roadblocks. Single-round *in vitro* transcription was performed using DNA templates encoding a fragment of the *E. coli* signal recognition particle (SRP) RNA with a template or nontemplate strand biotin modification in the absence and presence of SAv. The distribution of TECs does not change between 30 and 60 s of transcription. The percentage of complexes halted by the SAv roadblock at 60s is shown. **(B)** TECs are not dissociated by collision with a SAv roadblock. The DNA templates described in (A) were immobilized on SAv-coated magnetic particles to facilitate fractionation of single-round *in vitro* transcription reactions. RNAs within halted TECs are present in the pellet (P), whereas free RNAs are in the supernatant (S). The percentage of RNAs present in halted TECs is shown. Virtually all RNAs associated with halted TECs are partitioned into the pellet, indicating that TECs halted at SAv roadblocks are stable.

### Collision of RNAP with a streptavidin roadblock does not dissociate TECs

Because cotranscriptional SHAPE-Seq aims to probe RNAs in the context of a TEC, it is critical that collision of RNAP with a roadblock does not dissociate the TEC and release the RNA. To assess the stability of TECs stalled at a SAv roadblock, we immobilized each biotinylated template using SAv paramagnetic beads and separated RNAs in complex with a stable TEC from free RNAs by magnetic pull-down (Fig. 2B). We observed 95-98% of the RNAs in stalled TECs were still attached to beads after the pull-down, indicating that the vast majority of RNAs remain stably associated with RNAP in stalled TECs independent of the strand to which the SAv roadblock is tethered (Fig. 2B).

### Design of randomly biotinylated DNA templates for cotranscriptional SHAPE-Seq

Having established that SAv roadblocking can be used to halt RNAP in stable TECs, we next sought to validate its use in the cotranscriptional SHAPE-Seq experimental framework. Randomly biotinylated DNA templates were prepared by enzymatic incorporation of biotin-11-dNTPs during PCR amplification. Vent Exo- was selected for template amplification as it is particularly tolerant of biotin-11-dNTPs (28). To prevent biotinylation of the promoter nontemplate strand, which could interfere with promoter open complex formation, the forward PCR primer comprised positions -45 to -1 relative to the transcription start site (Fig. 3A). Optionally, the reverse primer can include a 5’ biotin modification as a terminal roadblock to prevent template run-off during the cotranscriptional SHAPE-Seq experiment (Fig. 3A). We prepared randomly biotinylated DNA templates for the *Bacillus cereus crcB* fluoride riboswitch (22) with targeted biotinylation levels of one, two, or four modifications per DNA template (Fig. 3B) and performed cotranscriptional SHAPE-Seq in the presence and absence of fluoride. This choice of model system allowed us to compare the general characteristics of the SAv roadblocking strategy with our previous Gln111 approach (10), and to perform a detailed comparison of the reactivities obtained from each method.

**Figure 3.**
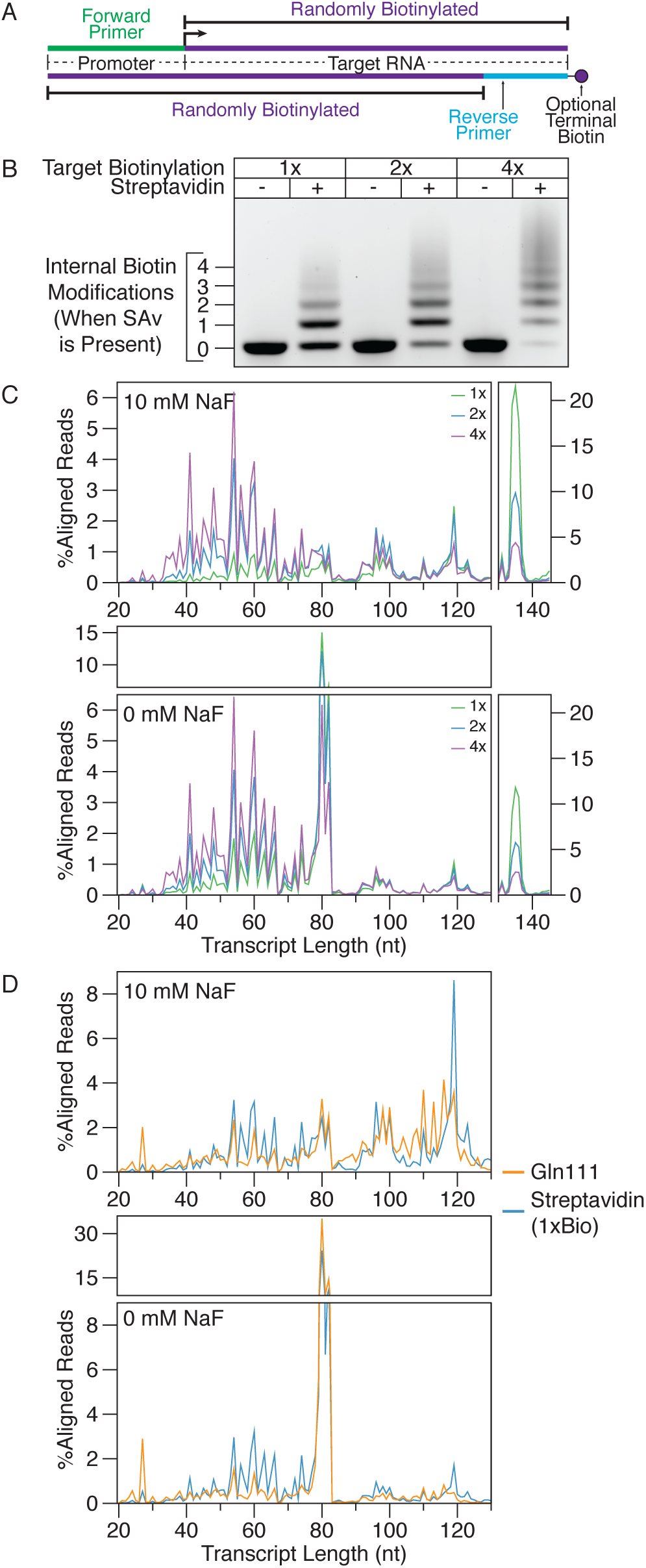
Distribution of aligned reads across transcript lengths. **(A)** Schematic of randomly biotinylated DNA templates used in cotranscriptional SHAPE-Seq experiments. **(B)** Gel shift assay showing the distribution in the number of internal biotins that are introduced during PCR with biotinylated dNTPs used at concentrations calculated to introduce an average of one, two, or four biotins per DNA template. **(C)** Distribution of read alignments over transcript lengths from cotranscriptional SHAPE-Seq with SAv roadblocking. Cotranscriptional SHAPE-Seq was performed on the *Bacillus cereus crcB* fluoride riboswitch with 10 mM or 0 mM fluoride using DNA templates with a targeted biotinylation rate of one, two, or four modifications per template. %Aligned Reads is calculated by dividing the unmodified RNA reads (SHAPE (−) sample) aligned at each transcript length by total unmodified reads aligned. ‘Full length’ terminally roadblocked TECs are separately plotted from internal stops to provide a clear view of the accumulation of both classes of stops. Fluoride riboswitch function is evident in the distribution of RNAs transcribed without fluoride, which accumulate as terminated products from position 80 - 82, compared to those transcribed with fluoride, which are distributed at positions beyond the terminator or accumulate at the terminal roadblock. **(D)** Comparison of read alignment distribution for SAv and Gln111 cotranscriptional SHAPE-Seq of the *Bacillus cereus crcB* fluoride riboswitch. Reads aligning to terminally roadblocked transcripts were not included when calculating the percentage of reads aligned to each transcript length since Gln111 roadblock run-through transcripts do not align due to the presence of an EcoRI site in the DNA template (10). Gln111 cotranscriptional SHAPE-Seq data downloaded from the Small Read Archive (http://www.ncbi.nlm.nih.gov/sra) (Table S1).

### Validation: Analysis of streptavidin and Gln111 transcript length alignments

To obtain a complete reactivity matrix for a target RNA, it is necessary to interrogate the structure of all intermediate transcripts. Thus, it is critical that all RNA intermediates are well represented in cotranscriptional SHAPE-Seq libraries. To assess coverage of the intermediate transcript lengths when biotin-SAv roadblocks are randomly incorporated, we examined the *Bacillus cereus crcB* fluoride riboswitch (22,29) with cotranscriptional SHAPE-Seq using varying degrees of biotinylation (1, 2, or 4 biotins/template) as described above. We then compared the distribution of unmodified transcript lengths to the number of biotins expected to be present in the DNA template (Fig. 3C). Increased DNA template biotinylation proportionally shifts the distribution toward shorter lengths as it becomes increasingly likely for RNAP to encounter a SAv roadblock (Fig. 3C). As an alternative to increased template biotinylation, enrichment for stalled complexes could also be achieved by omitting the terminal roadblock and using immobilized DNA templates to remove run-off transcripts before the transcription reaction is stopped (Fig. 2B).

In all samples the transcript lengths are distributed unevenly in a distinct pattern of peaks and troughs, representing high and low abundance, respectively. Interestingly, comparison of the alignment distributions produced by SAv and Gln111 roadblocking reveal remarkable consistency between both methods (Fig. 3D). One plausible explanation for the similarity of SAv and Gln111 transcript distributions is that the linker ligation step used to facilitate reverse transcription in the SHAPE-Seq v2.1 strategy (11) influences the representation of transcript lengths in cotranscriptional SHAPE-Seq libraries. There is well-documented structure- and sequence-dependent bias (30-32) in RNA-RNA ligation by T4 RNA ligase 2 truncated KQ (NEB). While such bias does not influence cotranscriptional SHAPE-Seq reactivity calculation, as reactivities are calculated internally for each length, reduction of ligation bias is a target of future development as it would reduce the sequencing depth necessary to adequately cover all intermediate transcript lengths by flattening the transcript length distributions. Because the transcript length distribution of cotranscriptional SHAPE-Seq libraries produced using SAv roadblocking approximates that of libraries produced using Gln111, we concluded that SAv roadblocking provides a sufficient distribution of TECs for reliable cotranscriptional SHAPE-Seq measurements.

### Validation: Analysis of streptavidin and Gln111 cotranscriptional SHAPE-Seq reactivity measurements

We next compared the cotranscriptional SHAPE-Seq reactivity measurements made for the *crcB* fluoride riboswitch using SAv roadblocks to previous measurements made using Gln111 roadblocks (10) (Table S2). Our previous characterization of the *crcB* fluoride riboswitch revealed key signatures of aptamer folding and fluoride binding as well as the fluoride-dependent bifurcation of the RNA folding pathway to produce the riboswitch ‘ON’ and ‘OFF’ regulatory states (10) (Fig. 4A). The same molecular signatures and their associated transitions are readily observable in reactivity matrices produced using SAv roadblocking (Fig. 4B-E, S1A-B, S2A-B) indicating that overall cotranscriptional SHAPE-Seq uncovers the same RNA structural information regardless of whether a SAv or Gln111 transcription roadblock is used.

We next compared SAv and Gln111 cotranscriptional SHAPE-Seq reactivities across all RNA intermediates by calculating reactivity differences (Δρ) (Fig. 4F-G, S1C-D, S2C-D). Interestingly, Δρ analysis reveals a notable distinction: when SAv roadblocking is used, the RNAP footprint can protect an additional upstream ~4 nt from SHAPE modification as can be seen by the presence of a stripe of higher Gln111 reactivities adjacent to the RNAP position in the Δρ matrices (Fig. 4F-G, S1C-D, S2C-D). Expanded protection of the nascent transcript from SHAPE modification can be explained by backtracking, the upstream movement of RNAP along the DNA template (33), when SAv roadblocking is used to halt TECs. This behavior is remarkably consistent, and is visible in all three SAv cotranscriptional SHAPE-Seq experiments, both in the presence and absence of fluoride (Fig. 4F-G, S1C-D, S2C-D).

**Figure 4.**
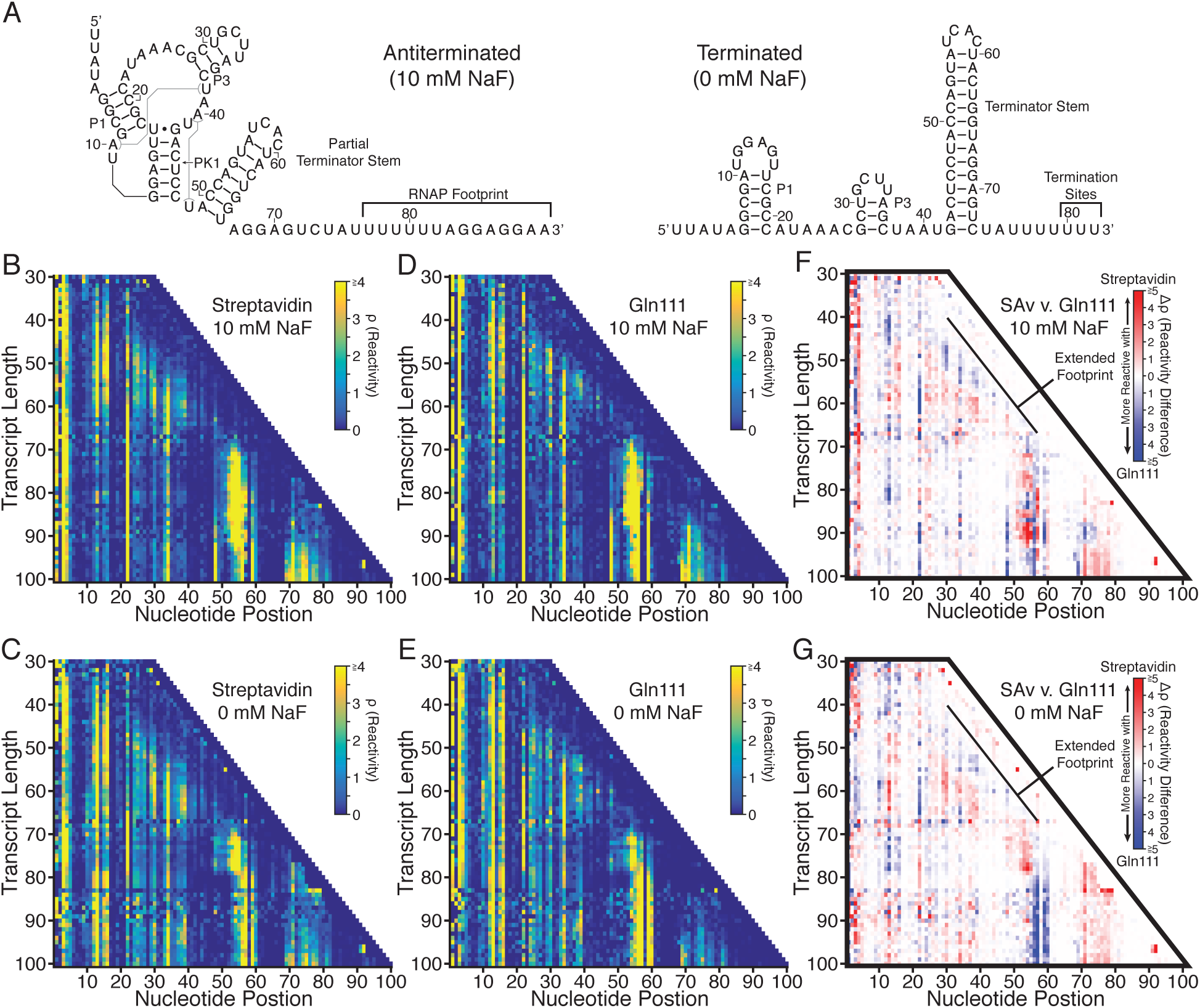
Cotranscriptional SHAPE-Seq of the *B. cereus crcB* fluoride riboswitch using SAv and Gln111 roadblocking. **(A)** Antiterminated and terminated secondary structures of the *B. cereus crcB* fluoride riboswitch. Structures are drawn according to phylogenetic (22), crystallographic (29), and cotranscriptional SHAPE-Seq (10) data. **(B-C)** Cotranscriptional SHAPE-Seq reactivity matrices produced using SAv roadblocking using 1xBiotin templates with 10 mM (B) or 0 mM (C) sodium fluoride (NaF). **(D-E)** Cotranscriptional SHAPE-Seq reactivity matrices produced using Gln111 roadblocking with 10 mM (D) or 0 mM (E) NaF (10). Gln111 data was downloaded from the RNA Mapping Database (RMDB) (http://rmdb.stanford.edu/repository/) (Table S2). **(F-G)** Reactivity differences (Δρ) between SAv and Gln111 roadblocking data with 10 mM NaF (F) and 0 mM NaF (G). Regions of low reactivity tend to have Δρ values that are close to 0, whereas regions with moderate or high reactivities exhibit greater Δρ values.

Consistent with the interpretation that the collision of RNAP with a SAv roadblock produces backtracked complexes, RNA folding transitions associated with aptamer folding and terminator nucleation are displaced downstream by 1-4 transcript lengths and appear to be more gradual when TECs are stalled with SAv (Fig. 5). The first such major structural transition that is observed earlier is the fluoride-independent decrease in P1 loop (nt 11 to 16) reactivity as it pairs with the lower 6 nt of the terminator stem (nt 42 to 47) to form the pseudoknot PK1 (Fig. 4A) (10) With Gln111, PK1 folds abruptly as P1 loop reactivity decreases sharply over transcript lengths 57 to 59, independent of fluoride concentration. With SAv, P1 loop reactivity decreases more gradually over transcript lengths 57 to 61, with an even more gradual drop in the absence of fluoride (Fig. 5A, S1E, S2E). Furthermore, reactivity changes at nucleotides A10 and A22 that were previously shown to be associated with aptamer folding (10) are also displaced downstream in the SAv dataset such that they remain coordinated with PK1 folding (Fig. S3).

**Figure 5.**
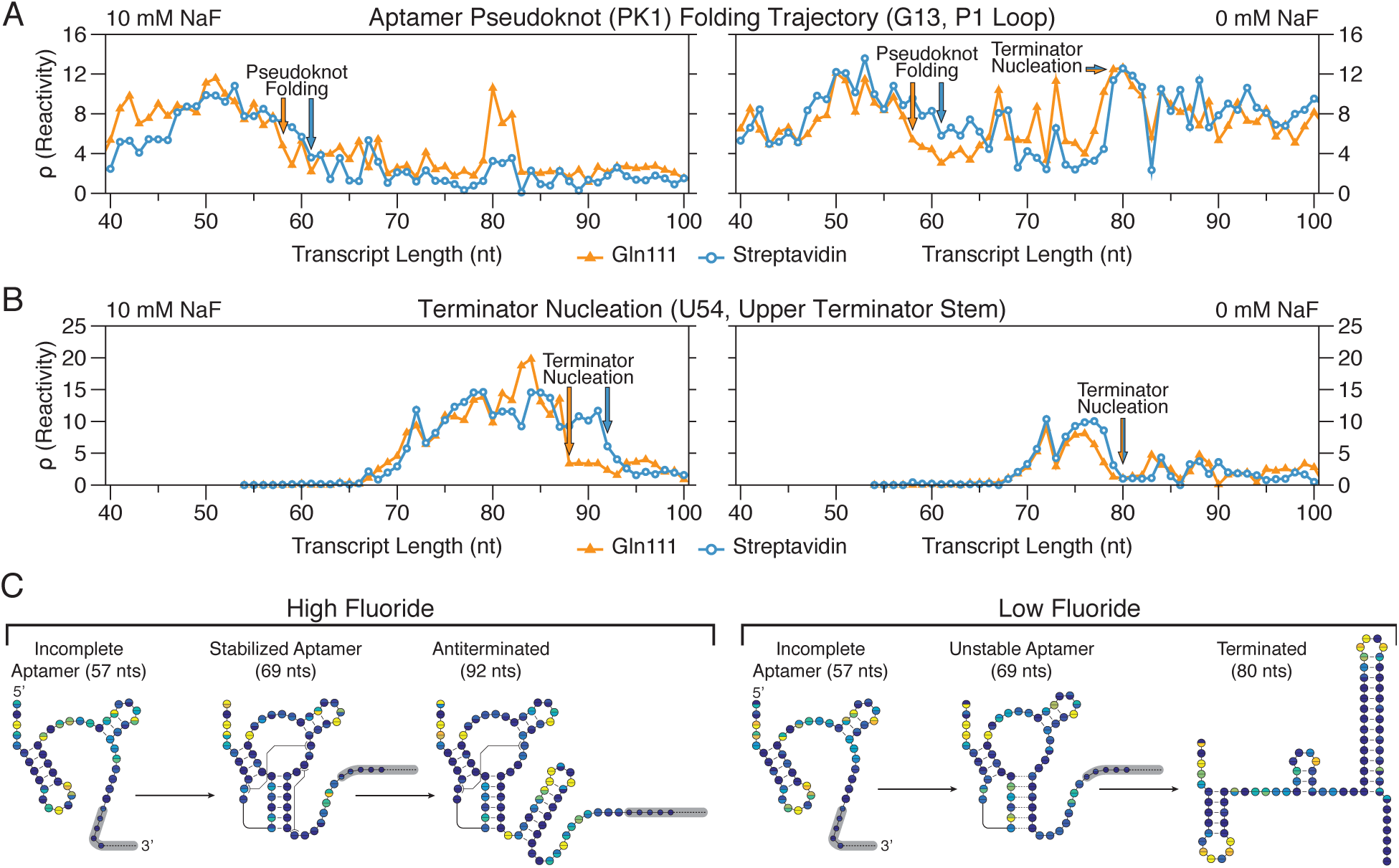
Comparison of *B. cereus crcB* fluoride riboswitch RNA folding transitions observed with SAv and Gln111 roadblocking. Gln111 data was downloaded from the RNA Mapping Database (RMDB) (http://rmdb.stanford.edu/repository/) (Table S2). **(A)** Cotranscriptional SHAPE-Seq reactivity trace of nucleotide G13 showing the fluoride-independent folding of pseudoknot PK1 followed by the fluoride-dependent bifurcation of the riboswitch folding pathway. **(B)** Cotranscriptional SHAPE-Seq reactivity trace of nucleotide U54 showing nucleation of the upper terminator stem. **(C)** *B. cereus crcB* fluoride riboswitch folding pathway. A model for the cotranscriptional folding pathway of the B. *cereus crcB* fluoride riboswitch in the presence (left) and absence (right) of NaF is shown. The transcript lengths shown are representative of key RNA structural states in the fluoride riboswitch folding pathway. Circles represent nucleotides and are colored according to cotranscriptional SHAPE-Seq reactivities obtained using SAv (top half of each circle) or Gln111 (bottom half of each circle) transcription roadblocks (Fig. 4B-E, Supplementary Figs. 4 and 5).

Following aptamer folding, the *crcB* fluoride riboswitch directs transcription termination or antitermination in the absence or presence of fluoride, respectively. In the absence of fluoride, terminator nucleation is observed as a coordinated reactivity decrease in the upper terminator stem (nt 52 to 55) and reactivity increase in the P1 loop as the terminator hairpin winds and disrupts PK1 (10). Terminator nucleation in the absence of fluoride is consistently displaced downstream by 1 transcript length when SAv roadblocking is used, occurring across lengths 76 to 79 with Gln111 and 77 to 80 with streptavidin (Fig. 5B, S1F, S2F). In the presence of fluoride, nucleation of the upper terminator stem is delayed until RNAP has traversed the poly-U tract and the fluoride-bound aptamer sequesters the base of the terminator stem so that only a partial terminator hairpin can form (10). In contrast to termination in the absence of fluoride, partial terminator nucleation in the presence of fluoride is more sensitive to the roadblock used, occurring at length 88 with Gln111 and length 92 with SAv (Fig. 5B, S1F, S2F). The relative insensitivity of terminator hairpin folding to roadblock type when nucleation occurs in close proximity to RNAP is consistent with observations that nascent RNA structure can prevent RNAP backtracking (34). Importantly, all RNA structural states associated with the *crcB* fluoride riboswitch termination and antitermination transitions are identified by cotranscriptional SHAPE-Seq regardless of the transcription roadblock used (Fig. S4).

The final noteworthy distinction between cotranscriptional SHAPE-Seq reactivities produced with SAv and Gln111 roadblocking is observed at the transcription termination sites in the presence of fluoride. Because *crcB* fluoride riboswitch antitermination efficiency in cotranscriptional SHAPE-Seq conditions is close to 100%(10), we do not expect to see reactivity signatures of transcription termination in the presence of fluoride. Nonetheless, previously we observed high P1 loop reactivity at the termination sites (80 to 82) when Gln111 is used to stall TECs (Fig. 5A). In contrast, P1 loop reactivity remains low at the termination sites when SAv roadblocking is used, suggesting that TECs stalled by SAv are resistant to this effect.

Our cotranscriptional SHAPE-Seq analysis of the *crcB* fluoride riboswitch with SAv and Gln111 roadblocking demonstrates that the choice of transcription roadblock can influence the transcript length at which a transition is observed by 1-4 nts. However, despite subtle distinctions in reactivity measurements, it is abundantly clear that both roadblocking strategies capture the same reactivity signatures associated with aptamer folding and the riboswitch regulatory decision (Figs. S4 and S5).

## Discussion

We have developed a sequence-independent method for distributing stalled TECs across a DNA template using SAv roadblocking and characterized the basic properties of SAv as an internal transcription roadblock. We also benchmarked the use of SAv roadblocks in cotranscriptional SHAPE-Seq against data that was previously generated using Gln111. We found that cotranscriptional SHAPE-Seq results were largely independent of the roadblock strategy used and propose that differences are dependent on the relative propensity for each transcription roadblock to induce backtracking. Indeed, the overall cotranscriptional SHAPE-Seq reactivity landscape of the fluoride riboswitch folding pathway is largely independent of the stalled TEC distribution strategy (Figs. 4B-E, S1A-B,S2A-B). We therefore suggest that while the use of Gln111 roadblocking for cotranscriptional SHAPE-Seq may provide greater accuracy than SAv roadblocking in mapping folding events to specific transcript lengths, the simplicity and reduced cost of SAv roadblocking makes it better suited for generation of full cotranscriptional SHAPE-Seq reactivity profiles.

The use of SAv roadblocking within the cotranscriptional SHAPE-Seq experimental framework both simplifies and reduces the cost of generating high-resolution profiles of RNA folding pathways. The use of SAv to halt TECs at all positions across a DNA template provides several advantages over Gln111 in the context of cotranscriptional SHAPE-Seq. First, amplification of DNA template libraries for SAv roadblocking only requires two primers, while Gln111 roadblocking requires a large primer set whose costs exceeds that of the biotin-11-dNTPs required for randomly biotinylated DNA template preparation, especially for long RNA targets. Second, whereas Gln111 is not commercially available, the use of SAv as a transcription roadblock requires only readily available reagents, thereby improving the accessibility of the method.

While streptavidin roadblocking provides several experimental advantages for cotranscriptional SHAPE-Seq, in some cases the use of Gln111 may prove advantageous because of its properties as a transcription roadblock. For example, while distributed streptavidin roadblocks provide a simple and effective method for elucidating full RNA folding pathways, Gln111 roadblocking is better suited for the interrogation of small subsets of intermediate transcripts. Perhaps the most intriguing property of Gln111 roadblocking in the context of cotranscriptional SHAPE-Seq is that it facilitates the identification and removal of reads from roadblock run-through products. Because the incorporation of an EcoRI site into a DNA template introduces non-native sequence, any TECs that ‘run-through’ the roadblock will synthesize an RNA containing aberrant sequence and can therefore be discarded during read alignment. In contrast, every DNA template used for streptavidin roadblocking contains the entirety of the target RNA sequence and therefore, reads generated by TECs that run-through the first roadblock they encounter and stop at a subsequent roadblock are indistinguishable from those that were successfully halted by the first roadblock encountered. This distinction does not influence cotranscriptional SHAPE-Seq results in the context of the *B. cereus crcB* fluoride riboswitch, but is an important consideration when studying RNAs that may be sensitive to transcription pausing because readthrough of a roadblock could function as a non-native pause.

The development of a second strategy for halting TECs across a DNA template afforded us the opportunity to examine the influence of transcription roadblock properties on cotranscriptional SHAPE-Seq data. The broad agreement of *B. cereus crcB* fluoride riboswitch reactivity matrices generated with each method is indicative of a high degree of experimental reproducibility, even across roadblocking strategies (Fig. 4, S5). Specifically, our analysis of the *B. cereus crcB* fluoride riboswitch revealed key molecular signatures depicting aptamer folding, fluoride binding, and the fluoride-dependent bifurcation of the riboswitch folding pathway into the terminated and antiterminated functional modes regardless of the roadblock used. The primary distinction between cotranscriptional SHAPE-seq measurements made using SAv and Gln111 roadblocking is that observed structural transitions are shifted upstream by 1-4 nts when SAv is used to halt TECs. This distinction suggests a degree of uncertainty in the location of RNAP relative to the 3’ end of the nascent transcript and is a direct consequence of using the RNA 3’ end to indicate RNAP location. While the RNA 3’ end generally provides a close approximation of the location of RNAP on a DNA template, it is not a direct measurement of RNAP position due to inherent variability of RNA 3’ end position relative to RNAP (35). Changes in the location of RNAP relative to the RNA 3’ end (36) by processes such as backtracking would sequester RNA proximal to the RNAP exit channel while simultaneously displacing the RNA 3’ end from the active center, resulting in an apparent downstream shift in structural transitions. Given the inherent flexibility of SAv roadblocks due to the necessity of tethering biotin to the DNA duplex by a linker, it is not surprising that RNAP position would be less defined following collision with SAv than following collision with the relatively static Gln111 roadblock. The use of biotin-dNTPs with a shorter linker could reduce the flexibility of the roadblock, however, to our knowledge only biotin-11-dNTPs have complete commercial availability. Given the complexity of interactions between RNAP and the nascent RNA, particularly in regard to the extent to which RNA folding can occur in the RNA exit channel (37-39), the stochastic nature of RNA folding is likely not conducive to precise assertions of specific nucleotides at which RNA folding events occur except in the most energetically favorable circumstances such as unhindered intrinsic terminator nucleation. Instead, the power of cotranscriptional SHAPE-Seq is in the high-resolution identification of coordinated structural rearrangements, such as those that arise from *crcB* fluoride riboswitch aptamer folding and subsequent fluoride binding (10). Analysis of *crcB* aptamer folding using SAv roadblocking clearly demonstrates that coordinated reactivity changes remain associated even when the transcript length to which they are mapped is shifted (Fig. S3).

Distributing TECs across all positions of a DNA template presents a technical challenge for which numerous solutions with unique advantages and disadvantages exist. Protein-based roadblocking strategies (9,18,20,21) are advantageous because they do not rely on the efficient incorporation of a modified nucleotide during transcription and do not require the incorporation of non-native nucleotides. Instead, protein roadblocks leverage the ubiquitous challenge of traversing a physical barrier along a DNA template and should therefore be generalizable to RNA polymerases beyond *E. coli* RNAP.

The availability of both SAv roadblocking and Gln111 roadblocking as strategies for halting TECs in cotranscriptional SHAPE-Seq allows users to tailor the method to specific experimental needs and expands the utility of a powerful RNA structure probing method.

## Methods

### DNA template preparation

DNA templates for *in vitro* radiolabeled transcription experiments were prepared by PCR amplification. 500 μl reactions included 411.25 μl H_2_O, 50 μl 10x ThermoPol Buffer (New England Biolabs), 6.25 μl 10 mM dNTPs, 12.5 μl 10 μM forward primer (Table S3), 12.5 μl 10 μM reverse primer (Table S3), 2.5 μl template plasmid DNA, and 5 μl of Vent Exo- DNA polymerase (New England Biolabs). 100 μl aliquots were amplified with a thermal cycling program consisting of 30 cycles using an annealing temperature of 55 °C. After thermal cycling, reactions were pooled into two 250 μl aliquots, mixed with 50 μl 3M sodium acetate (NaOAc) pH 5.5 and 1 mL 100% ethanol (EtOH) each, and stored at -80 °C for 30 min. After centrifugation, precipitated pellets were washed with 1.5 mL cold 70% EtOH, and dried using a SpeedVac. Dried pellets were pooled by dissolving in 30 μl H_2_O, fractionated by gel electrophoresis with a 1% agarose gel, and extracted using the QIAquick Gel Extraction Kit (Qiagen). Purified template was quantified using a Qubit Fluorometer (Life Technologies). Amplification of SRP DNA templates (Table S4) with a biotin modification at positions 33 and 42 relative to the transcription start site was directed with oligonucleotides A and B or C (Table S3), respectively.

To generate a DNA template containing a single nontemplate strand biotin modification, 10 μM oligonucleotide D and 10 μM oligonucleotide E (Table S3) in 50 μl of 1x ThermoPol buffer (New England Biolabs) was incubated at 95 °C for 5 min and annealed by incubating at 37 °C for 20 min. After chilling annealed oligonucleotides at 4 °C for 1 min, 10 U of ExoI was added and the sample was incubated at 37 °C for 30 min to remove excess oligonucleotides. The resulting DNA templates were first purified by using the QIAquick Purification Kit (QIAgen) and then run on a 1% agarose gel and gel extracted using the QIAquick Gel Extraction Kit (Qiagen). Purified DNA template concentration was measured using a Qubit fluorometer.

#### Amplification of Biotinylated DNA Templates

Randomly biotinylated DNA templates were prepared by PCR amplification and gel extraction as described above except that instead of supplying a dNTP mixture, each dNTP was added individually to a total of 100 nmol combined dNTP and biotin-11-dNTP. Assuming equal probability of incorporating a biotinylated or non-biotinylated dNTP, for 1x biotin incorporation, the nmol quantity of each biotin-11-dNTP included in the reaction was determined using the formula

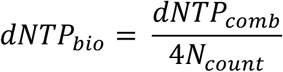

where *dNTP_bio_* is the nmol quantity of biotin-11-dNTP for base N included in the reaction, *N_count_* is number of occurrences of base N in the template and nontemplate strands of the DNA sequence that encodes the target RNA (not including reverse primer sequence), and *dNTP_comb_* is the combined nmol quantity biotinylated and non-biotinylated dNTP for base N included in the reaction. For higher biotin modifications, *dNTP_bio_* was then multiplied by the desired number of biotin modifications per template. The quantity of each non-biotinylated dNTP included in the reaction was determined by subtracting *dNTP_bio_* from *dNTP_comb_*. Biotin-11-dATP and biotin-11-dGTP were purchased from Perkin Elmer. Biotin-11-dCTP and biotin-11-dUTP were purchased from Biotium. Amplification of randomly biotinylated *B. cereus crcB* fluoride riboswitch DNA templates (Table S4) was directed by oligonucleotides F and G (Table S3).

### *In vitro* transcription (Radiolabeled)

For each sample, 0.125 pmol of biotinylated DNA template was pre-incubated with 12.5 pmol SAv monomer (Promega) at room temperature for 30 min. 25 μL reaction mixtures containing 5 nM DNA template/SAv complexes and 0.5 U NEB RNAP Holo (New England Biolabs) were incubated in transcription buffer (20 mM tris(hydroxymethyl)aminomethane hydrochloride (Tris-HCl) pH 8.0, 0.1 mM ethylenediaminetetraacetic acid (EDTA), 1 mM dithiothreitol (DTT) and 50 mM potassium chloride (KCl), 0.2 mg/mL bovine serum albumin (BSA), and 200 μM ATP, GTP, CTP and 50 μM UTP containing 0.5 μCi/μL [α-^32^P]-UTP for 10 min at 37 °C for 10 min at 37 °C to form open complexes. Single-round transcription was initiated by the addition of magnesium chloride (MgCl_2_) to 5 mM and rifampicin to 10 μg/ml. Transcription was stopped by adding 125 μl of stop solution (0.6 M Tris pH 8.0, 12 mM EDTA, 0.16 mg/mL tRNA).

For pellet/supernatant separation experiments, 10 μl of Streptavidin Magnesphere Paramagnetic Particles (Promega) per sample were first prepared by washing three times with SAv binding buffer (0.5 M sodium chloride (NaCl) and 20 mM Tris-HCl pH 7.5) before incubating with 0.125 pmol template DNA per sample in SAv binding buffer for 30 min. Bound templates were pulled-down and washed three times with 1x transcription buffer. *In vitro* transcription reactions were mixed as described above. After 30 s, reactions were placed on a magnetic stand and allowed to separate for 30 s before the supernatant was removed and added to 125 μl of stop solution and the pellet was resuspended in 125 μl of stop solution and 25 μl 1x transcription buffer.

Following *in vitro* transcription, RNAs were purified by the addition of 150 μL of phenol/chloroform/isoamyl alcohol (25:24:1), vortexing, centrifugation and collection of the aqueous phase. RNA was precipitated by adding 450 μl of 100% EtOH and storage at -20 °C. Following precipitation, RNA was resuspended in transcription loading dye (1x transcription buffer, 80% formamide, 0.05% bromophenol blue and xylene cyanol). RNAs were fractionated by electrophoresis using 12% denaturing polyacrylamide gels containing 7.5 M urea (National Diagnostics, UreaGel). Reactive bases were detected using an Amersham Biosciences Typhoon 9400 Variable Mode Imager. Quantification of bands was performed using ImageQuant. For all experiments, individual bands were normalized for incorporation of [α-32P]-UTP by dividing band intensity by the number of Us in the transcript. SAv roadblocking efficiency was calculated by dividing the sum of roadblocked RNAs by the sum of all roadblocked and run-off products. Aborted/paused products were not included in this calculation.

### In vitro transcription (Cotranscriptional SHAPE-Seq)

Reaction mixtures containing 100 nM randomly biotinylated DNA template and 4 U of *E. coli* RNAP holoenzyme (New England Biolabs) were incubated in transcription buffer, 0.2 mg/mL BSA, and 500 μM NTPs for 7.5 min at 37 °C to form open complexes. When present, NaF was included to a final concentration of 10 mM. Following open complex formation, SAv monomer (Promega) was added to 40 μM and incubated for another 7.5 min. Single-round transcription reactions were initiated by addition of MgCl_2_ to 5 mM and rifampicin to 10 μg/ml. After 30 seconds, cotranscriptional experiments were SHAPE modified by splitting the reaction and mixing half with 2.78 μL of 400 mM BzCN dissolved in anhydrous DMSO ((+) sample) or anhydrous DMSO only (Sigma Aldrich; (−) sample) for ~2s before addition of 75 μL of TRIzol solution (Life Technologies) and extraction. Extracted RNAs were dissolved in 20 μL 1× DNase I buffer (New England Biolabs) containing 1 U DNase I (New England Biolabs) and incubated at 37 °C for 30 min. After DNA digestion, 30 μL of H_2_O and 150 μL of TRIzol were added and the RNA was extracted and dissolved in 10 μL 10% DMSO.

### Sequencing Library Processing

An RNA linker was adenylated using the 5’ DNA adenylation kit (New England Biolabs), purified by TRIzol extraction, and quantified using a Qubit fluorometer as described in previously (10). Extracted RNAs were ligated to an RNA linker using T4 RNA Ligase 2 truncated KQ (New England Biolabs) by incubation at room temperatures described previously (10). Reverse transcription of the linker ligation products was performed using Superscript III Reverse Transcriptase (Life Technologies) as described previously (10). Ligation of an Illumina A_b adapter fragment was performed using CircLigase I ssDNA ligase (Epicentre) as described previously (10). ssDNA libraries were used to generate fluorescently labeled dsDNA libraries for library quality control as described previously (10). The resulting dsDNA libraries were analyzed by capillary electrophoresis using an ABI 3730xl and the resulting traces were used to evaluate library length distribution and the presence of adapter dimer prior to sequencing. Sequencing libraries were generated as described previously (11).

### Sequencing and Analysis

Sequencing was performed on the Illumina HiSeq2500 in Rapid Run mode using 2×36 bp paired end reads and 20% phiX. All cotranscriptional SHAPE-Seq computational tools used in this study can be found on GitHub at: https://github.com/LucksLab/Cotrans_SHAPE-Seq_Tools/releases/ and https://github.com/LucksLab/spats/releases/. Target FASTA files were prepared using the Cotrans_targets.py script as described previously (10). Reads were mapped and processed for Spats v1.0.1 as described previously (11).

## Acknowledgements

We thank the Lis Lab at Cornell University for the use of their facilities for radioactive materials experiments. This work was supported through a New Innovator Award through the National Institute of General Medical Sciences of the National Institutes of Health [grant number 1DP2GM110838 to JBL], and Searle Funds at The Chicago Community Trust [to JBL]. The content is solely the responsibility of the authors and does not necessarily represent the official views of the National Institutes of Health.

## Supplementary Materials

**Figure S1.**
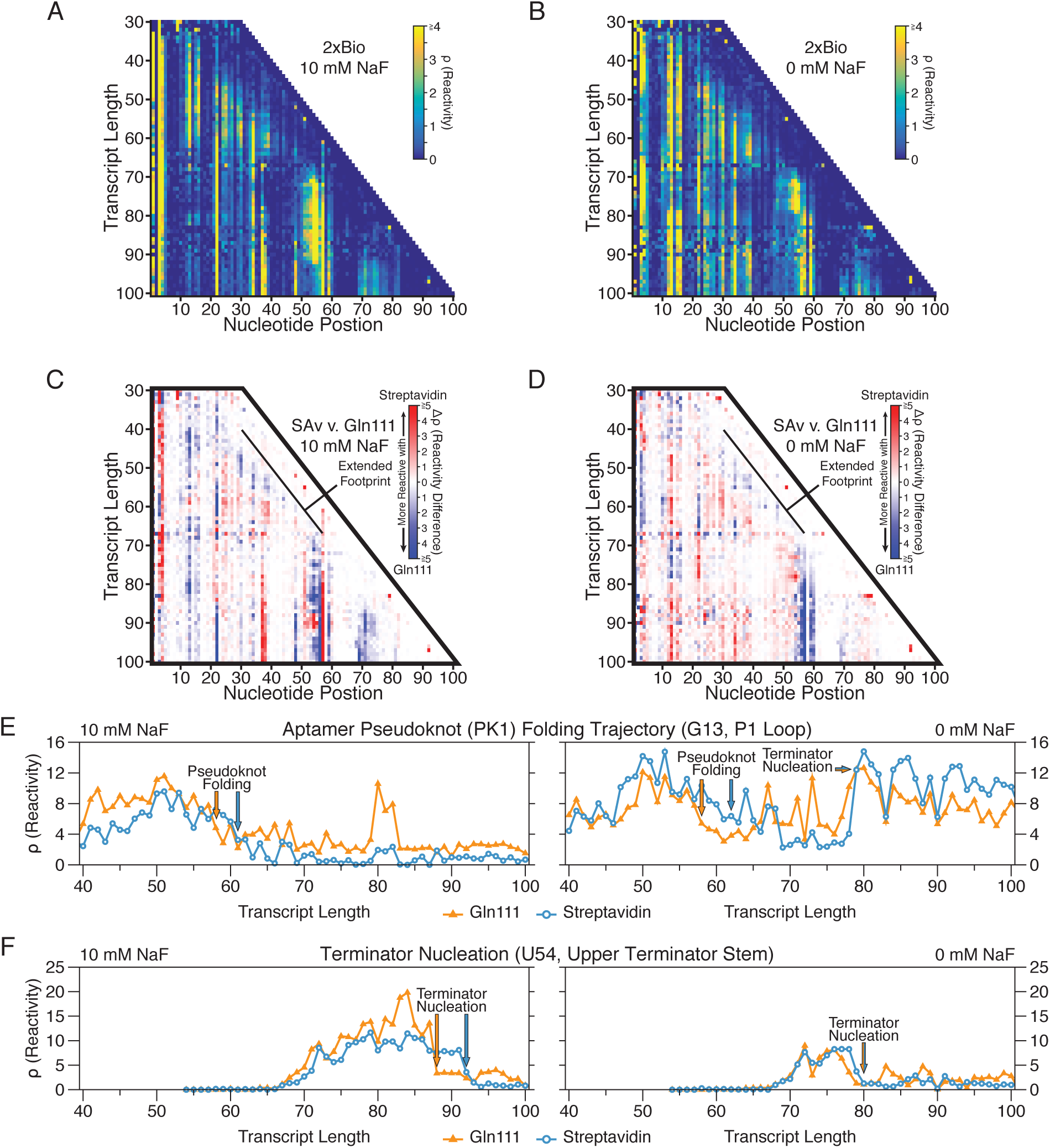
Comparing cotranscriptional SHAPE-Seq reactivities of the *B. cereus crcB* fluoride riboswitch using streptavidin (2x biotinylated template) with Gln111 roadblocking. Gln111 data was downloaded from the RNA Mapping Database (RMDB) (http://rmdb.stanford.edu/repository/) (Table S2). **(A-B)** Cotranscriptional SHAPE-Seq reactivity matrices produced using 2x streptavidin roadblocking with 10 mM (A) and 0 mM (B) NaF. **(C-D)** Reactivity differences (Δρ) between streptavidin and Gln111 roadblocking data with 10 mM (C) and 0 mM NaF(D). **(E)** Reactivity trace of nucleotide G13 showing the fluoride-independent folding of pseudoknot PK1 and the fluoride-dependent bifurcation of the riboswitch folding pathway. Key transitions are annotated. **(F)** Reactivity trace of nucleotide U54 showing nucleation of the upper terminator stem. Key transitions are annotated.

**Figure S2.**
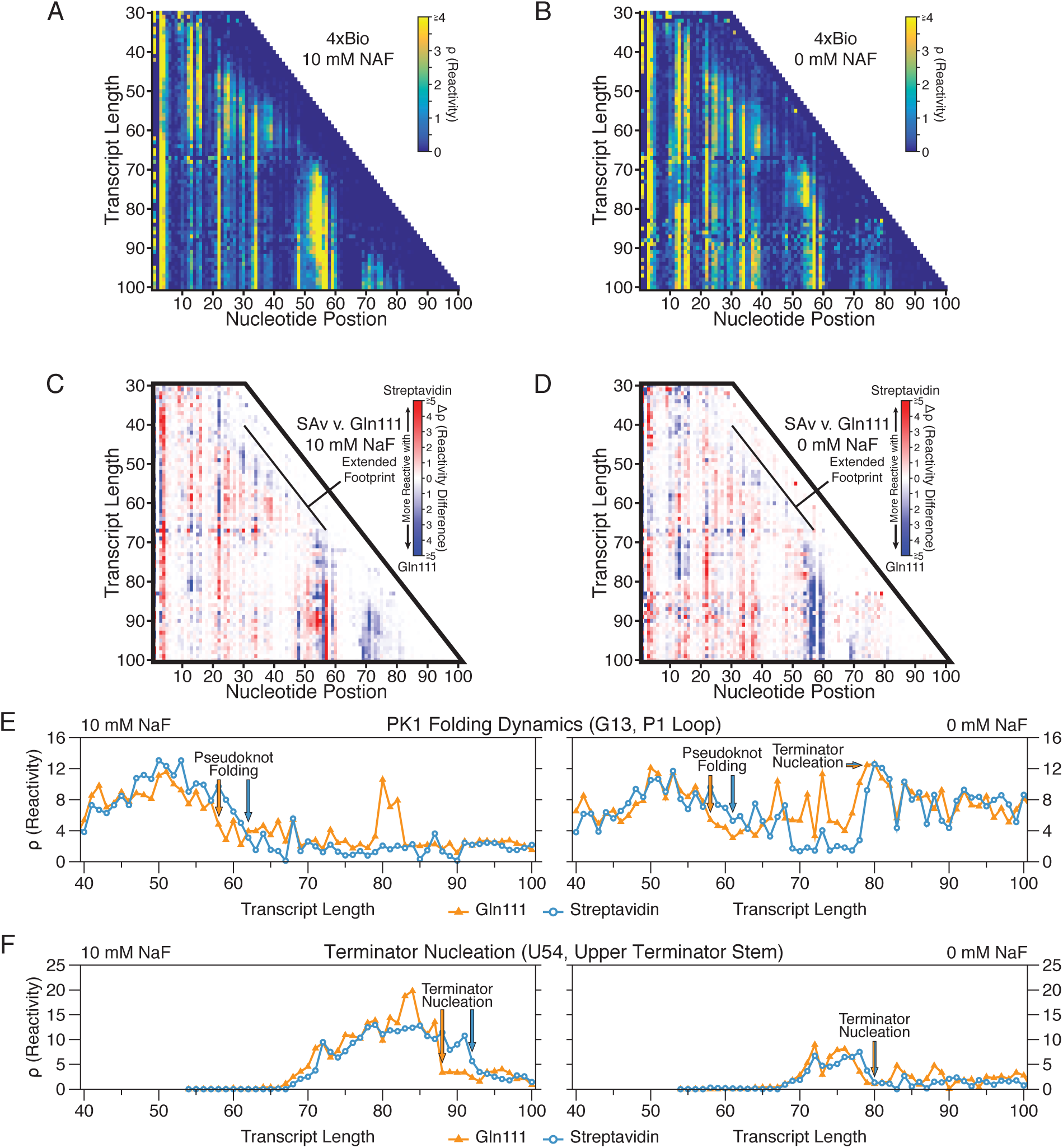
Comparing cotranscriptional SHAPE-Seq reactivities of the *B. cereus crcB* fluoride riboswitch using streptavidin (4x biotinylated template) with Gln111 roadblocking. Gln111 data was downloaded from the RNA Mapping Database (RMDB) (http://rmdb.stanford.edu/repository/) (Table S2). Data presented as in Fig. S1.

**Figure S3.**
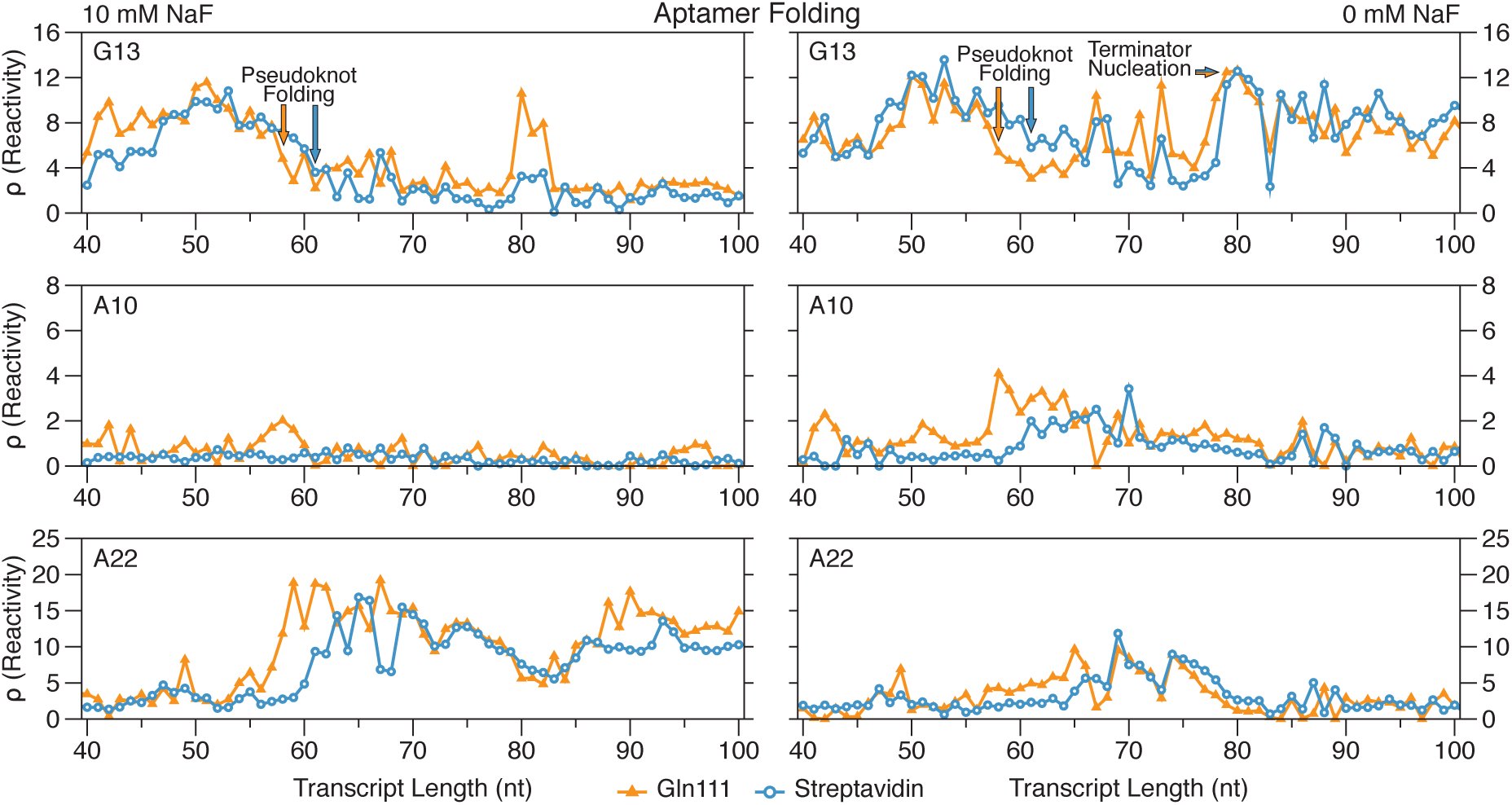
Coordinated displacement of aptamer folding reactivity transitions associated with streptavidin roadblocking. Streptavidin data (1× biotin) is from Figs. 4B and 4C. Gln111 data was downloaded from the RNA Mapping Database (RMDB) (http://rmdb.stanford.edu/repository/) (Table S2). Reactivity traces of nucleotides G13, A10, and A22 show the fluoride-independent folding of the *crcB* pseudoknot (G13), fluoride-dependent stabilization of the aptamer (A10, A22) and the fluoride-dependent bifurcation of the riboswitch folding pathway (G13). Reactivity changes at nucleotides A10 and A22 that are associated with pseudoknot formation and stabilization remain coordinated with aptamer folding regardless of the transcript length at which pseudoknot folding is observed.

**Figure S4.**
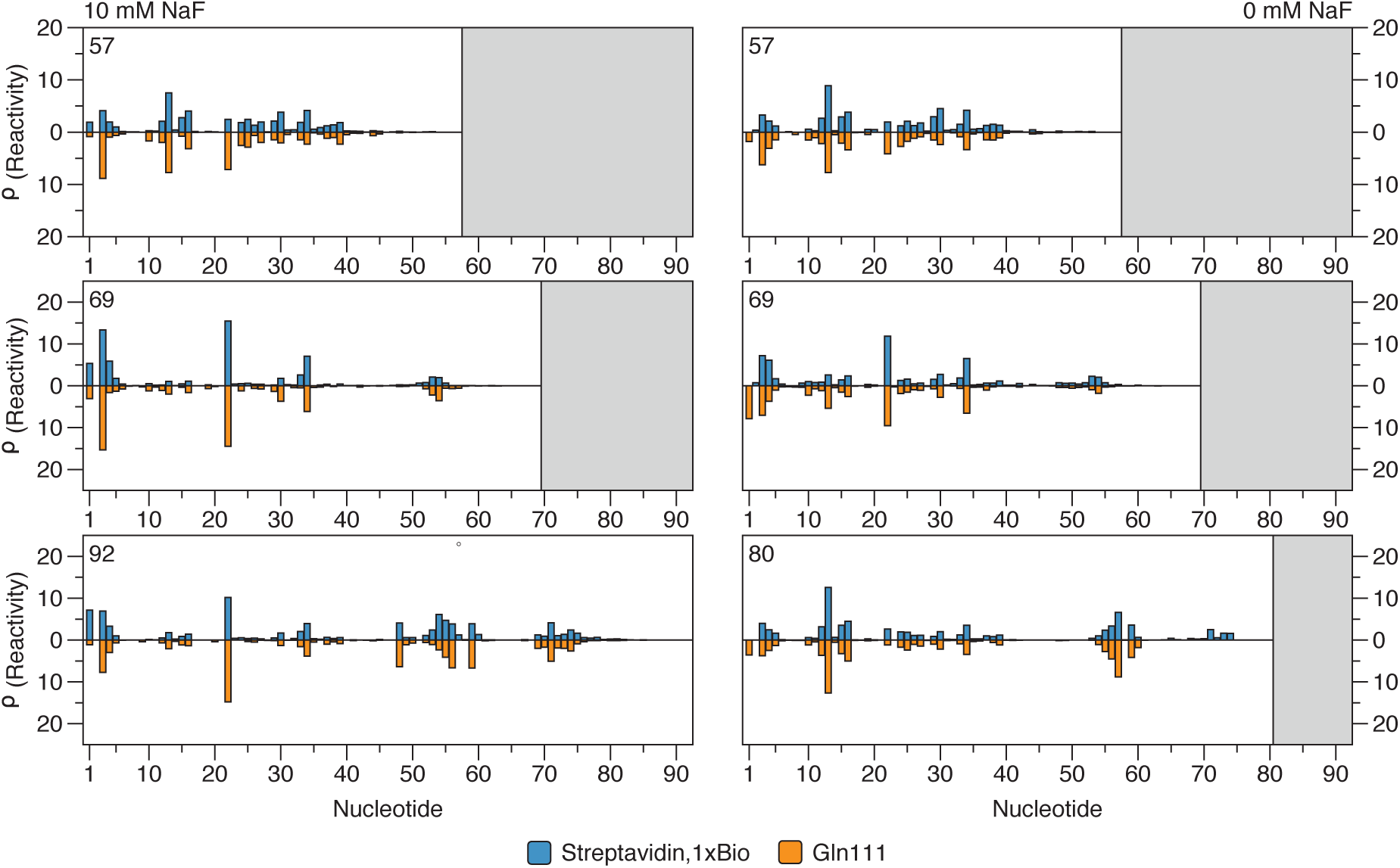
Cotranscriptional SHAPE-Seq identifies the RNA structural states associated with *crcB* aptamer folding with both streptavidin and Gln111 transcription roadblocks. Cotranscriptional SHAPE-Seq with 10 mM NaF reactivity profiles for transcript lengths 57, 69, and 92 (antiterminated) are shown. Cotranscriptional SHAPE-Seq with 0 mM NaF reactivity profiles for transcript lengths 57, 69, and 80 (terminated) are shown. Streptavidin data is taken from Figs. 4B and 4C. Gln111 data was downloaded from the RNA Mapping Database (RMDB) (http://rmdb.stanford.edu/repository/) (Table S2).

**Figure S5.**
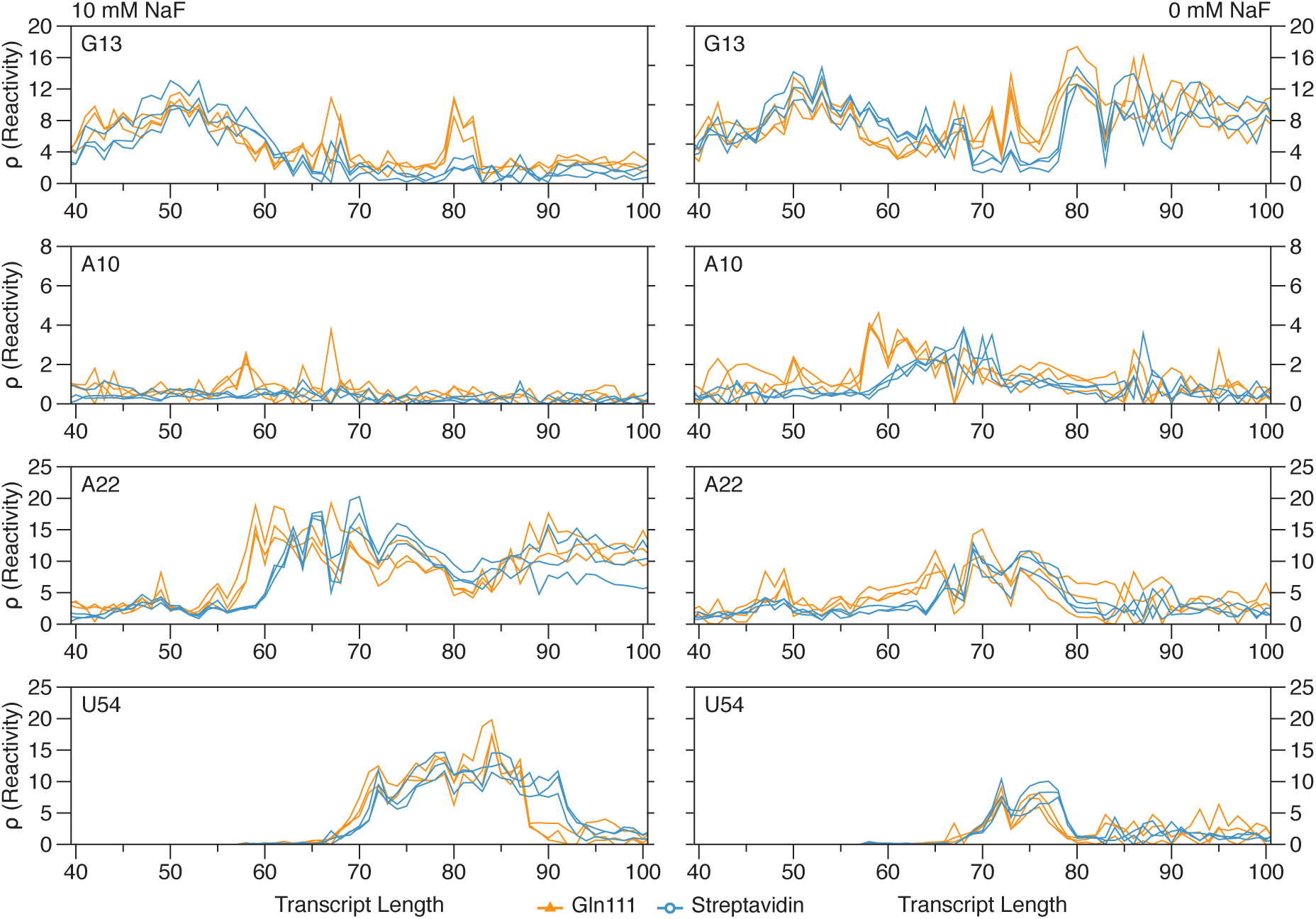
Reproducibility of cotranscriptional SHAPE-Seq reactivities with streptavidin and Gln111 transcription roadblocks. Streptavidin data is from Figs. 4B, 4C, S1A, S1B, S2A, and S2B. Gln111 data was downloaded from the RNA Mapping Database (RMDB) (http://rmdb.stanford.edu/repository/) (Table S2). Reactivity traces of nucleotide G13, A10, A22, and U54 are shown.

Sequencing data from Watters et al., 2016 used in this work were accessed through the Small Read Archive (http://www.ncbi.nlm.nih.gov/sra) BioProject accession number PRJNA342175. Individual BioSample Accession numbers are listed below:

**Table S1.** Small Read Archive (SRA) deposition table.

| **SRA Accession** | **RNA** | **Figure(s)** |
| --- | --- | --- |
| SRX2159297 | F- riboswitch, wt (replicate 1) | Fig. 3D |
| SRX2159317 | F- riboswitch, wt (replicate 1) | Fig. 3D |

Cotranscriptional SHAPE-Seq reactivity spectra from Watters et al., 2016 that were used in this work were accessed in the RNA Mapping Database (RMDB) (http://rmdb.stanford.edu/repository/), using the RMDB ID numbers indicated in the table below.

**Table S2.** RMDB data deposition table.

| **RMDB ID** | **RNA** | **Experiment** | **Figure(s)** |
| --- | --- | --- | --- |
| FLUORSW_BZCN_0001 | F- riboswitch, wt (replicate 1) | no ligand, cotranscriptional | Figs. 4, 5, S1, S2, S3, S4 & S5 |
| FLUORSW_BZCN_0002 | F- riboswitch, wt (replicate 2) | no ligand, cotranscriptional | Fig. S5 |
| FLUORSW_BZCN_0003 | F- riboswitch, wt (replicate 3) | no ligand, cotranscriptional | Fig. S5 |
| FLUORSW_BZCN_0004 | F- riboswitch, wt (replicate 1) | 10 mM ligand, cotranscriptional | Figs. 4, 5, S1, S2, S3, S4,& S5 |
| FLUORSW_BZCN_0005 | F- riboswitch, wt (replicate 2) | 10 mM ligand, cotranscriptional | Fig. S5 |
| FLUORSW_BZCN_0006 | F- riboswitch, wt (replicate 3) | 10 mM ligand, cotranscriptional | Fig. S5 |

Below is a table of oligonucleotides used for the preparation of *in vitro* transcription DNA templates. Abbreviations within primer sequences are as follows: ‘/iBiodT’ is a biotin modified thymidine residue with a linker length of 11, ‘/5Biosg/’ is a 5’ biotin moiety. These abbreviations were used for compatibility with the Integrated DNA Technologies ordering notation.

**Table S3.**
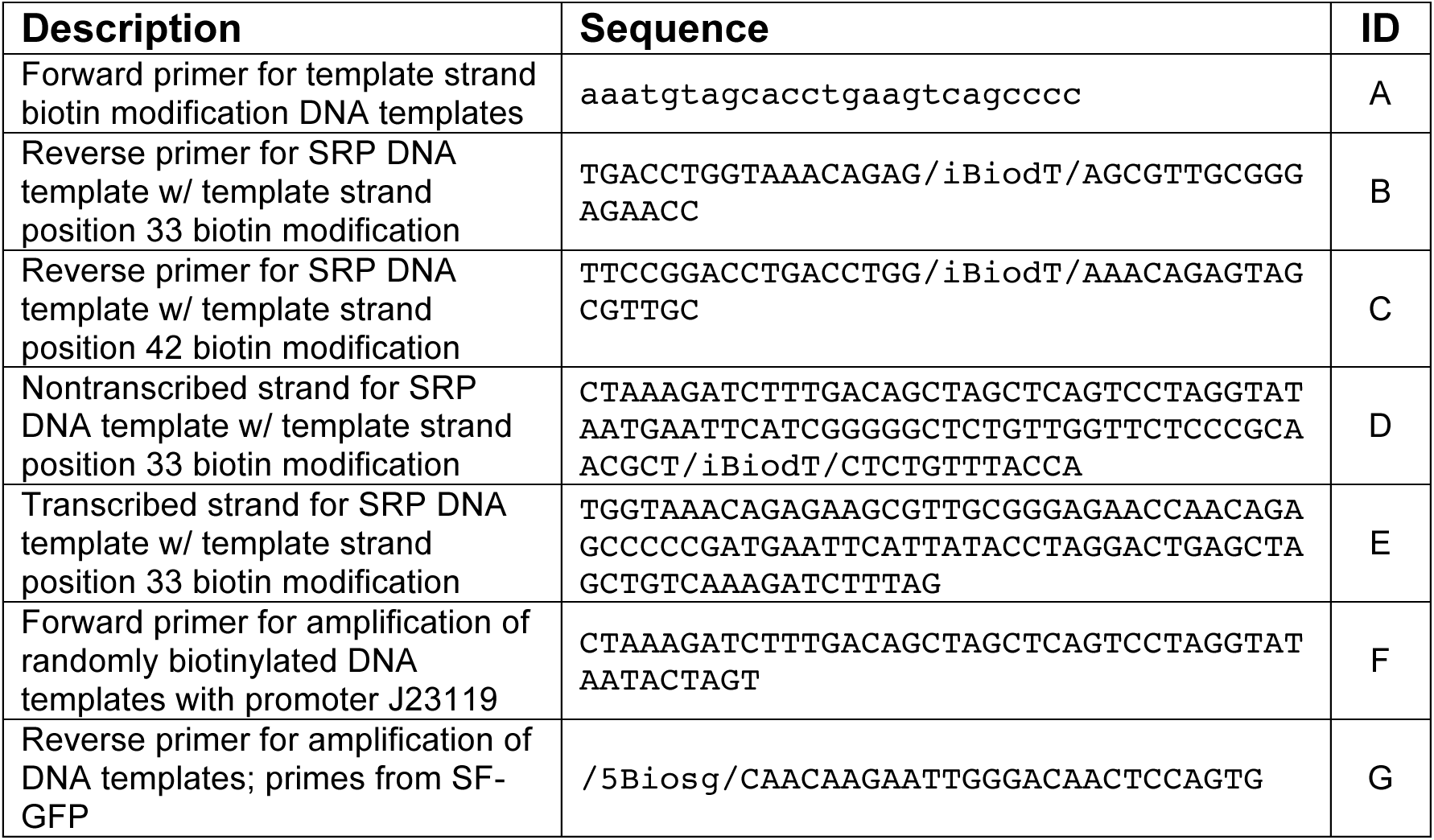
Oligonucleotides used for DNA template amplification or assembly.

Sequences of DNA templates used for *in vitro* transcription. Signal Recognition Particle (SRP) RNA sequence is as described in Wong et al. (6). A ribosome binding site (RBS) and superfolder GFP (SFGFP) sequence was included downstream of the *B. cereus crcB* fluoride riboswitch. Promoter sequences are blue. Mutation positions are highlighted with red.

**Table S4.**
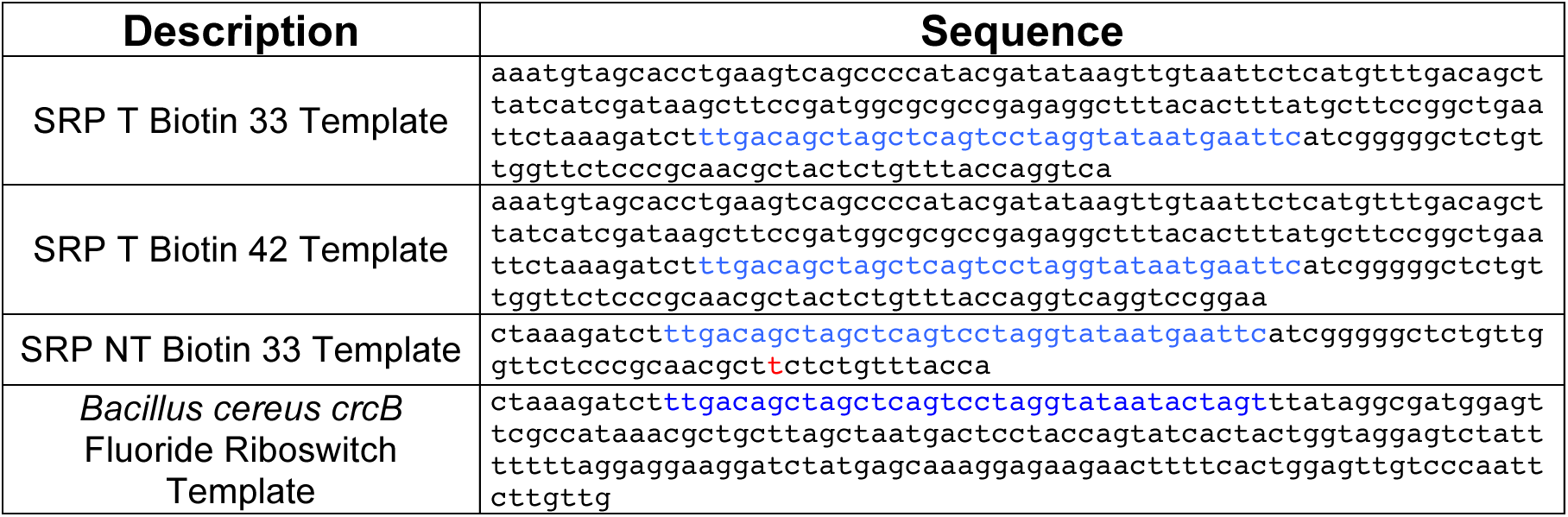
DNA templates used for in vitro transcription.

